# Human colon-on-a-chip enables continuous *in vitro* analysis of colon mucus layer accumulation and physiology

**DOI:** 10.1101/740423

**Authors:** Alexandra Sontheimer-Phelps, David B. Chou, Alessio Tovaglieri, Thomas C. Ferrante, Taylor Duckworth, Cicely Fadel, Viktoras Frismantas, Sasan Jalili-Firoozinezhad, Magdalena Kasendra, Eric Stas, James C. Weaver, Camilla A. Richmond, Oren Levy, Rachelle Prantil-Baun, David T. Breault, Donald E. Ingber

## Abstract

**Background & Aims:** The mucus layer in the human colon protects against commensal bacteria and pathogens, and defects in its unique bilayered structure contribute to intestinal disorders, such as ulcerative colitis. However, our understanding of colon physiology is limited by the lack of *in vitro* models that replicate human colonic mucus layer structure and function. Here, we investigated if combining organ-on-a-chip and organoid technologies can be leveraged to develop a human-relevant *in vitro* model of colon mucus physiology.

**Methods:** A human colon-on-a-chip (Colon Chip) microfluidic device lined by primary patient-derived colonic epithelial cells was used to recapitulate mucus bilayer formation, and to visualize mucus accumulation in living cultures non-invasively.

**Results:** The Colon Chip supports spontaneous goblet cell differentiation and accumulation of a mucus bilayer with impenetrable and penetrable layers, and a thickness similar to that observed in human colon, while maintaining a subpopulation of proliferative epithelial cells. Live imaging of the mucus layer formation on-chip revealed that stimulation of the colonic epithelium with prostaglandin E2, which is elevated during inflammation, causes rapid mucus volume expansion via an NKCC1 ion channel-dependent increase in its hydration state, but no increase in *de novo* mucus secretion.

**Conclusion:** This study is the first to demonstrate production of colonic mucus with a physiologically relevant bilayer structure *in vitro*, which can be analyzed in real-time non-invasively. The Colon Chip may offer a new preclinical tool to analyze the role of mucus in human intestinal homeostasis as well as diseases, such as ulcerative colitis and cancer.

---

The mucus layer in the human colon normally protects the intestinal epithelial cells against enormous numbers of luminal commensal bacteria and potential pathogens present in the gut lumen of healthy individuals ^1–5^. The human colonic mucus layer has a unique bilayer structure as it is composed of an inner layer that is normally impermeable to bacteria and a permeable outer layer ^1, 2, 6^. The integrity of the inner layer is most crucial in preventing direct contact of the bacteria with the colonic epithelium and associated chronic inflammation ^1–4^. In addition, changes of mucus layer homeostasis can indirectly influence intestinal barrier function^6^. Increased direct contact between these bacteria and the colonic epithelium can lead to gut barrier dysfunction and bacterial penetration through the epithelial tissue boundary. This can trigger injury and inflammation, for example, the mucus layer becomes penetrable to bacteria in dextran sodium sulfate-treated mice that develop colitis long before infiltration of immune cells is observed ^4, 5, 7^. Importantly, the inner colonic mucus layer in patients with ulcerative colitis (UC), a common form of chronic inflammatory bowel disease affecting the colon ^8^, also has been shown to allow bacterial penetration ^5, 6^. Prostaglandin E2 (PGE2) is elevated during intestinal inflammation, as in patients with UC ^9^, where it plays an essential role in wound healing ^9–11^. Recently, PGE2 also has been shown to be a direct mediator of fluid secretion using intestinal epithelial organoids ^12^. This function is thought to be mainly mediated through activation of ion channels leading to ion flux-driven water flux into the lumen ^12^. In the past, short term PGE2 treatment also has been reported to induce mucus secretion in cAMP-dependent manner mediated thought activation of its receptor EP4 in murine intestinal loop studies and mouse proximal colon explants ^13–15^. But the effect of PGE2 on the colonic mucus layer height and properties have been controversial and are not fully understood ^13–16^.

Unfortunately, it is impossible to study intestinal mucus physiology and changes in its behavior over time within the lumen of the living human colon to address these types of questions. Mucus physiology can be studied *in vivo* in animal models (e.g., using intestinal loop studies) ^13, 14^; however, these methods are highly invasive, technically challenging, and often only low resolution imaging is possible ^17^. More importantly, there are species-specific differences in mucus layer thickness and microstructure ^1, 2, 6^, and thus, there has been a search for new methods that could advance investigation in this area ^6, 17^.

Common challenges using *in vitro* cell culture models to investigate intestinal mucus physiology include the use of cancer-derived epithelial cells, such as Caco-2 and HT29-MTX cells, which results in secretion of the gastric mucin MUC5AC, but not typical intestinal MUC2 ^17, 18^. Primary human intestinal organoids can produce mucus, but because it is entrapped in the central lumen of the organoid, it is difficult to investigate its physiology ^17^. A few studies have demonstrated accumulation of mucins on the surface of primary human ileal or rectal organoid fragments cultured on Transwell (TW) inserts. But these cultures only accumulate thin (< 36 μm thick) mucus layers ^19^, which are significantly thinner than those observed *in vivo* (200–700 μm). Importantly, neither these cultures nor any other experimental *in vitro* method can reproduce the physiologically important bilayer structure that is seen in human colonic mucus ^17^. Thus, most studies on mucus biology rely on the use of short-lived *ex vivo* mouse or human tissue explants ^20^. While most of our current understanding of the colon mucus layer originated from studies of tissue explants ^4, 6, 15, 20, 21^, they have significant limitations including the need for repeated access to clinical biopsy specimens. In addition, clinical samples can only be used once, they cannot be used for long-term (> 1 day) cultures, and because the cells cannot be expanded *in vitro*, it is difficult to replicate results using the same donor ^17^.

Human organ-on-a-chip (Organ Chip) microfluidic culture technology has been used to create a human Small Intestine Chip ^22, 23^ and Colon Chip ^24^, which are respectively lined by primary human organoid-derived duodenal or colonic intestinal epithelial cells, offers an alternative approach to confront this challenge. Organ Chips offer many advantages over the intestinal organoids from which they are derived, including the ability to continuously collect samples from both the luminal and abluminal compartments; the Intestine Chip also exhibits greater transcriptomic similarity to *in vivo* intestine compared to organoids ^22^. In the present study, we set out to explore whether Organ Chip technology could be used to develop a method to recapitulate human colonic mucus layer formation *in vitro*.

## RESULTS

### Reconstitution of a polarized colonic epithelium

Primary colonic epithelial cells isolated from human colon resections and endoscopic biopsies (Supplementary Figure 1A) were first grown as colon organoids (Supplementary Figure 1B), which were then fragmented and cultured on top of a porous extracellular matrix-coated membrane with 7 μm diameter pores in the top channel of a 2-channel microfluidic Organ Chip device composed of optically clear poly-dimethyl siloxane (PDMS), as previously described ^22, 24^ (Figure 1A, Supplementary Figure 2A). Hank’s balanced salt solution (HBSS) and stem cell expansion medium were continuously perfused (60 ul h^-1^) through the top (epithelial) and bottom channels, respectively. When cultured under these dynamic flow conditions, the human colonic epithelial cells formed a confluent flat monolayer by day 3 that developed into a dense undulating epithelial sheet lined by columnar colonic cells by day 7 (Figure 1B, Supplementary Figure 2B-D). The colonic epithelium retained this dense morphology for at least 2 weeks in culture (Supplementary Figure 2B-D) and covered the entire channel uniformly (Supplementary Figure 2C).

**Figure 1:**
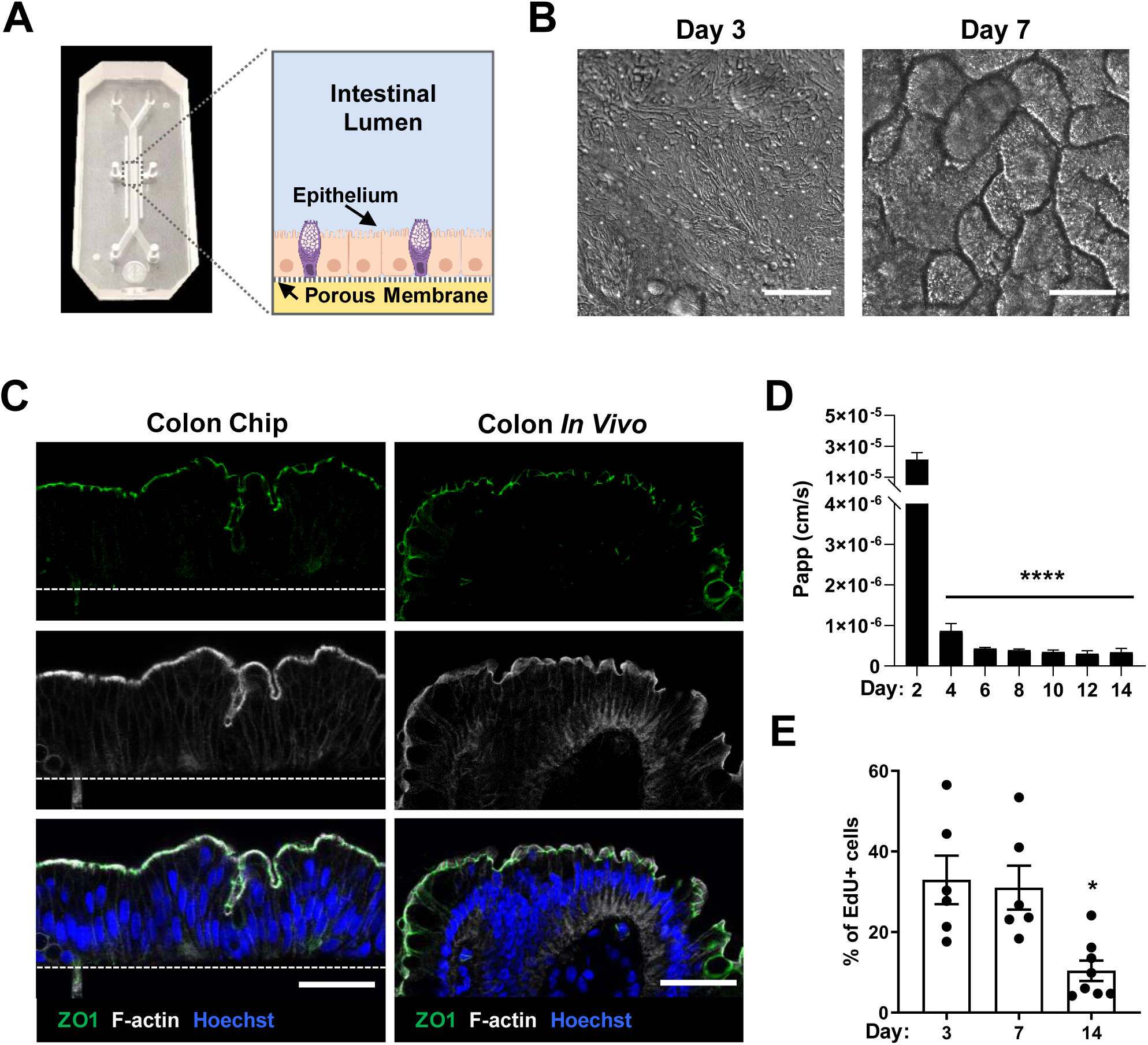
Patient-derived human Colon Chip. (A) Photograph and schematic cross-sectional view of the primary human Colon Chip microfluidic device with the colonic epithelium cultured in the top channel on the upper surface of the porous membrane that separates it from the bottom channel. (B) DIC images of the colonic epithelium 3 and 7 days after monolayer formation in the Colon Chip (viewed from above; bar, 100 µm). (C) Immunofluorescence confocal microscopic images of histological cross sections of the Colon Chip compared with human colon *in vivo* showing a polarized epithelium with tight junctions labeled with ZO1 (green) and brush border stained for F-actin (gray) restricted to the apical regions, and Hoechst-stained nuclei (blue) localized at the cell base (bar, 50 µm; white dashed line, top of the porous PDMS membrane in the Colon Chip). (D) Intestinal barrier function of the colonic epithelium measured over 14 days of culture on-chip by quantifying the apparent permeability (P_app_) of Cascade blue (550 Da) (n=3-11 chips; **** p*<0.0001* compared to day 2). (E) Quantification of cell growth by measuring EdU incorporation over 18 h using flow cytometry (n=6-8 chips, 2 experiments compiled; \**p*<0.05 compared to d3 and d7). All data represent mean ± SEM.

**Figure 2:**
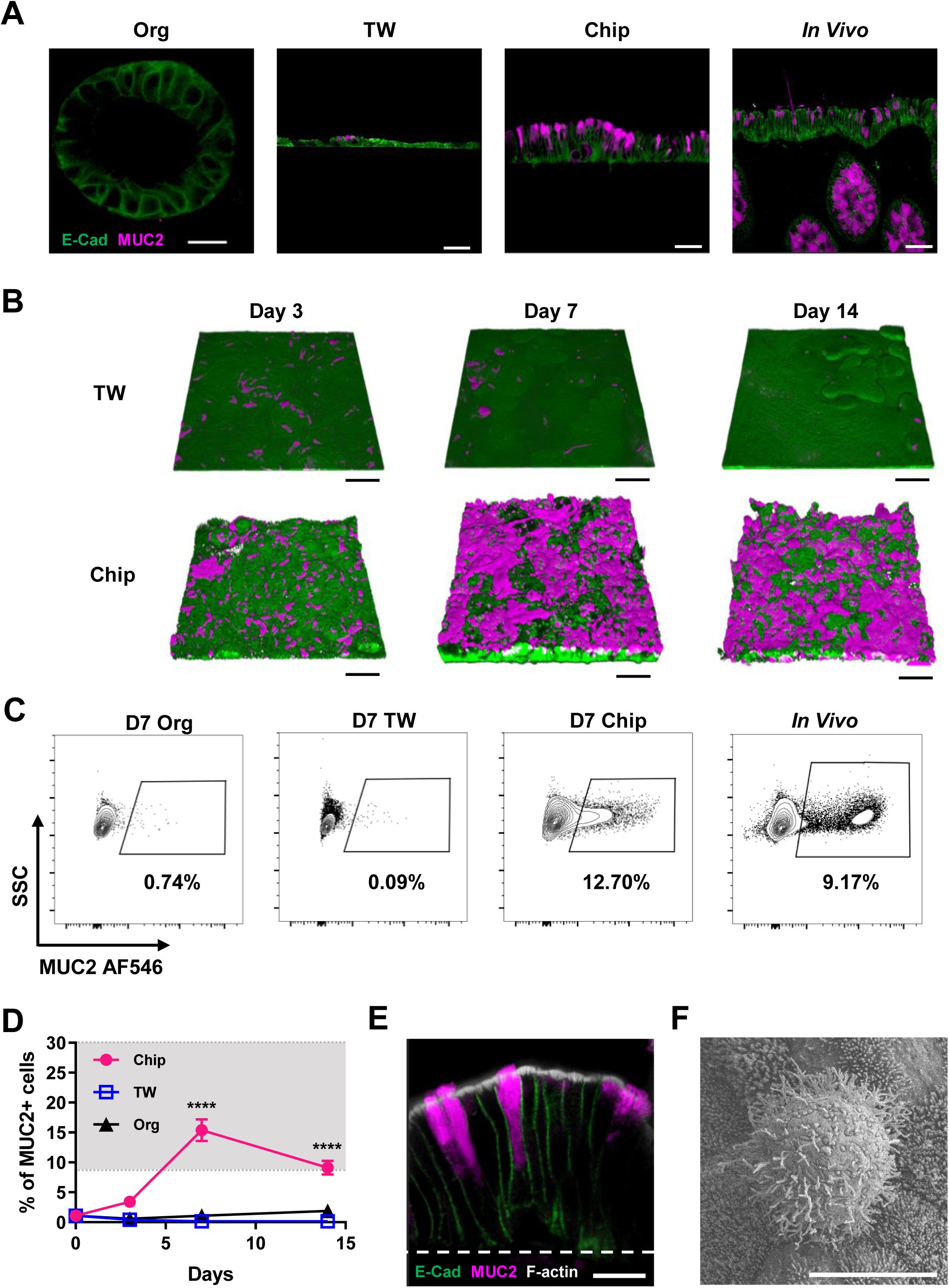
Spontaneous goblet cell differentiation in the Colon Chip. (A) Cross-sectional, confocal immunofluorescence microscopic images of epithelial cells in a colon organoid (Org) (bar, 20 µm), TW culture, Colon Chip (Chip), and human colon tissue section (all bars, 50 µm) stained for E-Cadherin (green) and MUC2 (magenta); cultures shown are 7 days after monolayer formation. (B) 3D reconstructions of confocal immunofluorescence microscopic z-stack images of epithelial cells in TW cultures and Colon Chips at day 3, 7 and 14 after monolayer formation showing epithelium stained for E-Cadherin (green) and MUC2 (magenta) (bars, 100 µm). (C) Contour plots of flow cytometric quantification of MUC2+ cells in colon organoid (Org), TW culture and Colon Chip (Chip) at day 7 of culture versus cells isolated from human colon tissue (*In Vivo*) and a graph (**d**) showing the full analysis at days 3, 7, and 14 (gray area indicates approximate range of *in vivo* values; n=3-9 devices, 2 experiments compiled; **** *p*<0.0001 compared to TW and Org; data represent mean ± SEM). (E) Higher magnification view of an immunofluorescence confocal micrograph showing cross section of the epithelium within the Colon Chip containing goblet cells stained for MUC2 (magenta), E-cadherin (green), and F-actin (gray) (bar, 20 µm). (F) Scanning electron micrographic image of the apical surface of a goblet cell within the epithelium cultured in the Colon Chip surrounded by enterocytes with apical microvilli (bar, 5 µm).

The polarized structure of the colonic epithelium is crucial for intestinal barrier function, as well as for uptake and transport of ions and water in the colon^25^. Immunofluorescence confocal microscopy confirmed that the colonic cells formed a polarized columnar epithelium in the Colon Chip at day 7 that closely resembled that seen in human colon *in vivo*, as evidenced by restriction of nuclei to the basal cytoplasm and appearance of ZO-1-containing tight junctions and an F-actin-rich brush border along the apical surface of the epithelium (Figure 1C), as well as basolateral localization of E-cadherin-containing adherens junctions (Supplementary Figure 3). Analysis of barrier function measured with the small fluorescent tracer, Cascade blue (550 Da), confirmed that the colonic epithelium formed a tight intestinal barrier beginning at 4 days of culture on-chip, which was maintained over 2 weeks (Figure 1D), as previously observed in a human Duodenum Chip ^22^. Importantly, under these culture conditions, the human colonic epithelium also retained a subpopulation of proliferating cells during the entire 2 weeks of culture (Figure 1E).

**Figure 3:**
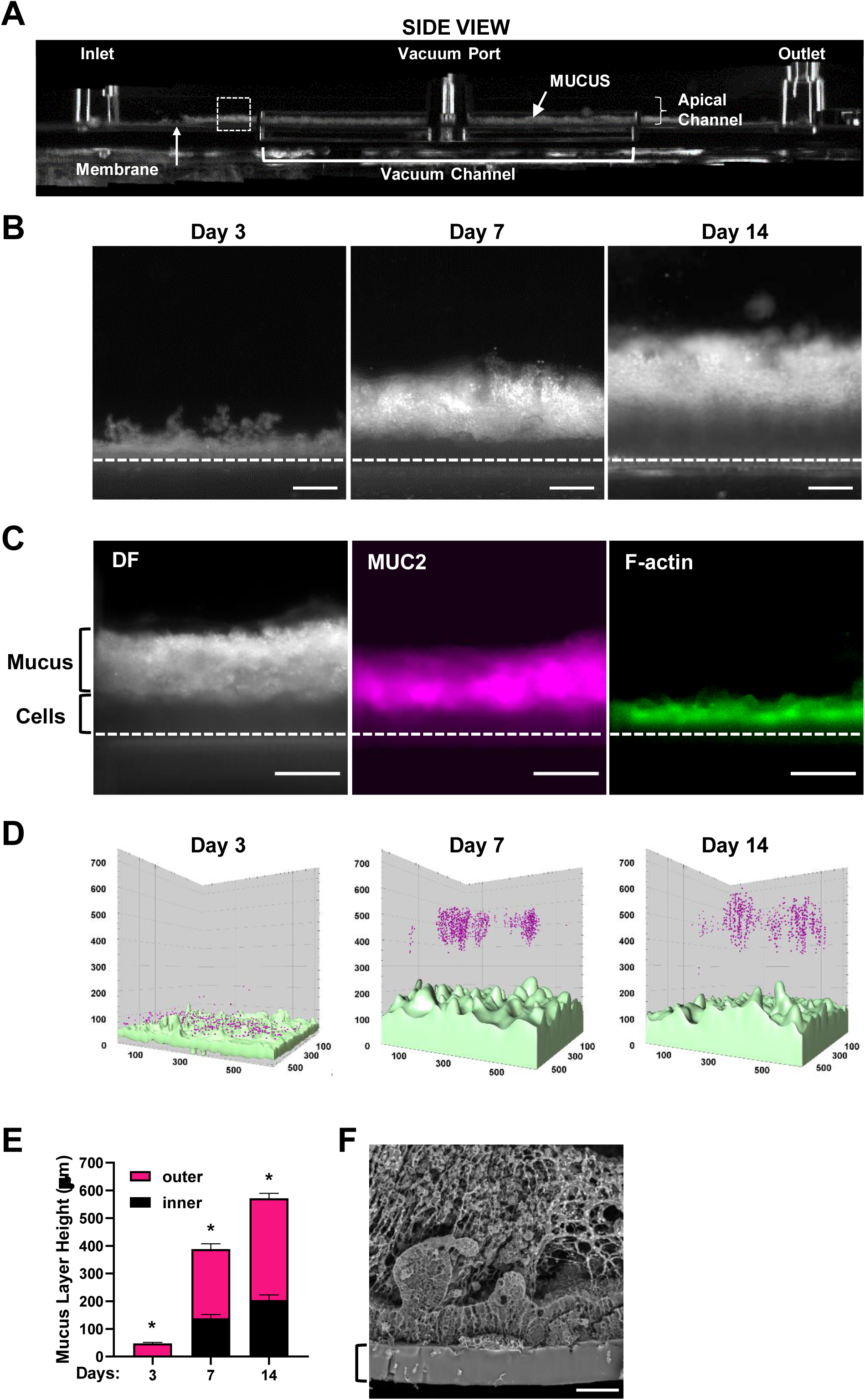
Formation of a mucus bilayer in the human Colon Chip. (A) Dark Field (DF) microscopic image of the side of an optically clear Colon Chip showing the entire length of the device with horizontal microfluidic upper and lower channels along its length, as well as upper inlet, outlet and vacuum port; a side vacuum channel is also apparent in this view. Note that mucus can be detected in the bottom half of the apical channel above the epithelium, which is cultured on the horizontal membrane that separates the upper and lower channels (dashed square corresponds to region shown in B and C). (B) Live DF images of the same Colon Chip at days 3, 7 and 14 after monolayer formation showing a reflective mucus layer that progressively increases in thickness over time above the darker cell region (dashed line, upper surface of the porous membrane; bar, 200 µm). (C) Higher magnification images of a region similar to the area shown in the dashed square in **a** within a Colon Chip fixed at day 7, demonstrating the presence of a thick mucus layer visualized by DF microscopy and MUC2 staining (MUC2) overlying the F-actin-rich brush border of the colonic epithelium (F-actin) (dashed line, upper surface of the porous membrane; bars, 200 µm). (D) A pseudo color 3D reconstruction of representative z-stack confocal images of the Colon Chip perfused with 1 µm fluorescent beads (magenta) 3, 7 or 14 days after monolayer formation; the cells were live stained with Calcein AM (light green). (E) Quantification of the thickness of the inner (impenetrable) and outer (penetrable) mucus layers base on the distribution of the beads (n = 3 chips, 3 independent regions per chip; all values significantly different between all days, * *p*<0.05 for inner and outer layer; data represent mean ± SEM). (F) Scanning electron micrograph of a cross section of the epithelium within a Colon Chip demonstrating the presence of a filamentous network within the mucus layer in direct contact with the apical surface of the cells (bracket indicates the PDMS membrane; bar, 50 µm).

### Goblet cell differentiation

The presence of goblet cells in the colonic epithelium is a critical requirement for any study of mucus physiology as these are the specialized intestinal cells that produce and secrete mucin 2 (MUC2), which is a major component of intestinal mucus ^26^. MUC2 polymers are densely packed in large secretory vesicles in goblet cells, which give the cells their typical “goblet” shape ^26^. As expected based on past work that showed stem cell expansion medium drives the proliferation of stem cells in organoid cultures ^27^, we found that our organoids, and TW cultures created using cells isolated from these organoids, formed few, if any, goblet cells at 1 week of culture when cultured in this medium (Figure 2A), and similar results were obtained even when TW cultures were maintained for 14 days (Figure 2B). Surprisingly, however, when the same organoid-derived colonic epithelial cells were cultured in the Colon Chip in the same stem cell expansion medium, high levels of goblet cell differentiation were observed, as indicated by appearance of MUC2+ epithelial cells that were similar in morphology and number to those seen in histological sections of human colon (Figure 2A). These MUC2+ cells could be detected as early as 3 days of culture and they expanded greatly in number by 1 to 2 weeks covering larger areas of the epithelium (Figure 2B).

Flow cytometric analysis of live epithelial cell populations harvested from colon organoids, TWs, Colon Chips, and human colonic tissue confirmed that there was little goblet cell differentiation in the TW and organoid cultures, whereas ∼15% of the epithelial cells differentiated into goblet cell in the Colon Chip, which similar to the percentage of goblet cells we measured in human colon tissue samples (∼10-30% depending on the donor; Figure 2C,D, Supplementary Figure 4A). Furthermore, the MUC2+ cells on the Colon Chip exhibited the typical goblet cell shape with apical location of mucus granules as observed *in vivo* (Figure 2E) ^26^, as well as a similar surface morphology when analyzed by scanning electron microscopy (Figure 2F) ^28^.

**Figure 4:**
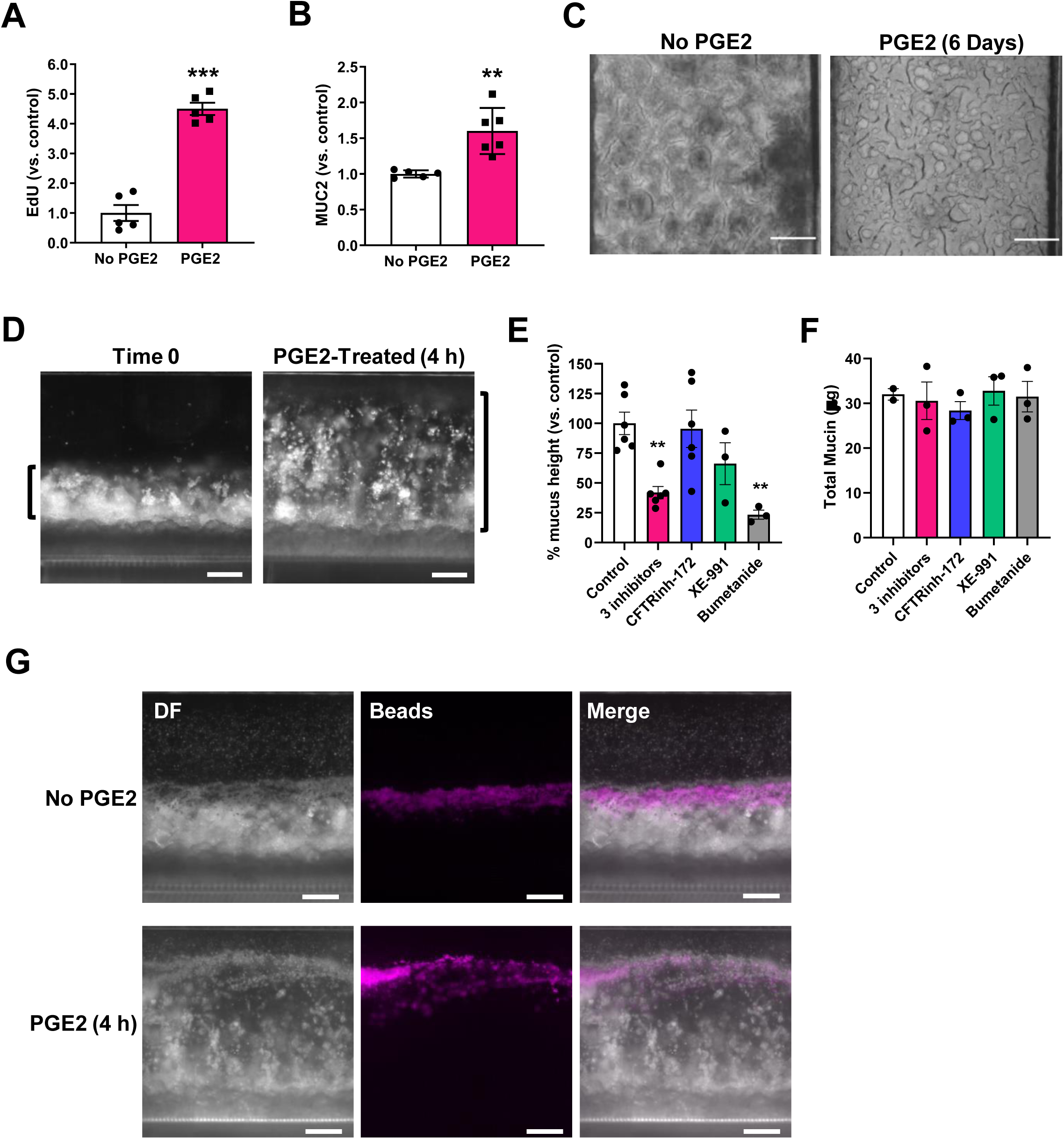
PGE2-induced mucus layer swelling on-chip. Graphs showing changes in EdU incorporation measured over 18 hours (A) and MUC2+ cells (B) in epithelial monolayers within Colon Chips after 6 days of PGE2 treatment relative to untreated controls (No PGE2) detected using flow cytometry (n = 5 chips for A; n = 5-6 chips for B; 2 experiments compiled). (C) Bright field images from above of Colon Chips that were untreated (No PGE2) or treated with PGE2 for 6 days (bars, 200 µm). (D) DF side view images of a Colon Chip with epithelial cells and mucus layer before and after PGE2 treatment for 4 hours (bars, 200 µm). Quantification of changes in mucus layer height based on the DF images (E) and total mucin production measured using an alcian blue assay (F) induced by PGE2 alone (control) or PGE2 in the presence of CFTRinh-172 (50 µM), XE-991 dihydrochloride (20 µM), bumetanide (100 µM) or a combination of all three ion channel inhibitors (data are presented relative to control chips treated with PGE2 alone (n = 2-6, 2 experiments compiled). (G) Side view images of Colon Chip with or without 4 h PGE2 treatment with 1 µm fluorescent beads (magenta) overlaid on DF images (bars, 200 µm). (** *p*< 0.01, **** *p*< 0.0001 vs. control; all data represent mean ± SEM).

Importantly, despite supporting spontaneous goblet cell differentiation, the Colon Chip cultures were simultaneously able to maintain a proliferative cell subpopulation at levels similar to those present in the organoid and TW cultures (Supplementary Figure 4B). While past work has shown that some goblet cell differentiation can be induced in TW cultures and organoids by replacing the stem cell expansion medium with differentiation medium ^27, 29, 30^, these cultures cease proliferating and are short-lived. Thus, the Colon Chip effectively provided an environment for goblet cell differentiation under conditions in which proliferative cells also remain present as occurs in human colon *in vivo*, while other culture models do not.

### Analysis of intestinal mucus accumulation and bilayer structure in living cultures

Given the spontaneous differentiation of large numbers of goblet cells in the Colon Chip that produce MUC2, which is the main mucin in colonic mucus, we next investigated if a physiologically relevant mucus bilayer forms on-chip. The existence of a mucus layer within the lumen of the apical epithelial channel was suggested by the appearance of increasing opacity of the Colon Chip over time when viewed from above by light microscopy (Supplementary Figure 2B, C). Importantly, because the microfluidic Organ Chip is optically clear and has defined linear channel geometries (Figure 1A), we were able to develop a method to visualize live cultures in cross section across the entire channel. This was accomplished by slicing ∼ 2 mm of the PDMS material away from the sides of the upper and lower channels, then turning the device 90° on its side and analyzing it using Dark Field (DF) microscopy (Figure 3A-C, Supplementary Figure 5A, B). Using this method, we detected the formation of a highly scattering layer above the cells that was well-developed by 1 week and when we analyzed the same chips over time, it continued to increase in height over the following week (Figure 3B). This layer stained positively for MUC2 and resided on top of the apical F-actin-rich brush border of the epithelium (Figure 3C), confirming that this material that accumulated over time in the Colon Chip is indeed a thick mucus layer. Fixation of intestine tissue with paraformaldehyde has been shown to produce a compression artifact (thinning of the layer) and disrupt the inner layer ^17, 31^; thus, an important advantage of this method is that it enables the *in situ* study of mucus development in its native state over an extended period of time in living cultures.

One of the unique properties of human colonic mucus is its bilayer structure, which is characterized by an inner layer that is impenetrable by bacteria and an outer penetrable mucus layer that is normally populated by commensal microbes. This feature of the mucus layer has been previously probed in tissue explants utilizing fluorescent 1 µm diameter microbeads to mimic bacteria ^20^. When we flowed similar fluorescent microbeads through the lumen of the epithelial channel and allowed them to settle, we observed a bead-free region approximately 200 µm in height above the apical membrane of the epithelial cells at days 7 and 14 (Figure 3D, E), which is similar to the thickness of the dense impenetrable inner mucus layer observed in human colon tissue explants using this method ^20^. The spread of the beads trapped in the mucus above the impenetrable layer corresponded to the penetrable outer mucus layer (Figure 3D, E). When we quantified the total height of the mucus layer (both inner and outer layers) formed on chip, we found that it reached ∼570 µm by day 14 (Figure 3E). Scanning electron microscopic analysis of a cross-section of the Colon Chip also revealed the presence of a fibrous network within the thick mucus layer in tight contact with the upper surface of the epithelium (Figure 3F), which again is similar to what has been observed in tissue explants analyzed using lectin staining ^32^. Thus, the microfluidic human Colon Chip is the first *in vitro* method to support spontaneous accumulation of a mucus layer in tight apposition to the apical surface of a cultured human colonic epithelium that replicates the thickness and unique bilayer properties displayed by human colonic mucus *in vivo*.

### Modeling the response of colonic epithelium to PGE2 on-chip

Prostaglandin E2 (PGE2) has been reported to be elevated in patients with ulcerative colitis, and it appears to contribute to healing of intestinal ulcers by increasing cell proliferation and altering mucus physiology ^9–11, 13–15^. To explore if the human Colon Chip method can be used to study this response *in vitro*, we perfused PGE2 (1.4 nM) through the basal channel from day 2 to 8. This 6-day exposure to PGE2 resulted in a 4.5-fold increase in proliferating cells compared to control chips, as measured by EdU incorporation (Figure 4A), as well as a 1.6-fold increase in MUC2+ cells (Figure 4B). The rise in EdU incorporation is consistent with reports that PGE2 promotes cell proliferation and is essential for wound healing ^9–11^.

In these studies, we noticed that the PGE2-treated chips developed blockage of the apical channel preventing fluid flow, suggesting that there also might be increased mucus production. This was surprising because the PGE2-treated Colon Chips appeared to contain less light-obscuring material compared to control Chips when imaged from above by bright field microscopy (Figure 4C), which stands in direct contrast to the increased opacity associated with the mucus layer development we observed in our earlier studies. Past animal and tissue explant studies exploring the effect of PGE2 on mucus production have produced contradictory results. Short term PGE2 treatment has been reported to increase colonic mucus accumulation in murine intestinal loop studies and mouse proximal colon explants ^13–15^, but no increase in mucus secretion could be detected in response to PGE2 treatment in isolated human colonic crypts ^16^ or distal colon mouse tissue explants^15^.

Thus, we set out to evaluate the short-term effect of PGE2 on mucus layer height using the human Colon Chip. Short term (4 h) treatment of the Colon Chip with PGE2 resulted in an increase in mucus height by 321.8 ± 44.3 µm, which is equivalent to a ∼2-fold increase compared to control chips (Figure 4D, Supplementary Figure 6A). Interestingly, however, there did not appear to be an associated increase in the total amount of mucins when quantified using an alcian blue-binding assay (Supplementary Figure 6B). This result suggests that the increase in height we detected was not due to secretion of more mucin materials, but rather to swelling of the pre-existing mucus.

While PGE2 has a multitude of functions in intestinal physiology, it plays an important role in the regulation of fluid secretion via ion transport ^12^. Because mucus production has been shown to depend on fluid secretion in mouse small intestine ^33, 34^, we tested if this property of PGE2 was responsible for its ability to rapidly increase mucus height. PGE2-induced fluid secretion can be blocked by inhibiting ion channels ^12^, and so we pretreated the human Colon Chip with a combination of 3 different ion channel inhibitors, CFTRinh-172, XE-991, and bumetanide, which are chemical inhibitors of the cystic fibrosis transmembrane conductance regulator (CFTR), K_V_7 (KCNQ) voltage-gated potassium channel, and NKCC1 Na-K-Cl cotransporter, respectively, before infusing PGE2. The combination of the 3 different ion channel inhibitors led to a significant reduction in the PGE2-induced increase in mucus height confirming the importance of ion transport in this hydration response (Figure 4E, Supplementary Figure 7).

To analyze the effects of each ion channel inhibitor individually, different Colon Chips were pre-treated with each of these ion channel inhibitors before exposing them to PGE2. Suppression of the CFTR and Kv7 K+ channel activity had no significant effect, however, inhibition of the basolateral NKCC1 Na-K-Cl cotransporter with bumetanide significantly reduced PGE2-induced mucus layer height on-chip (Figure 4E, Supplementary Figure 7). None of these channel inhibitors altered total mucin content (Figure 4F). Thus, the increase in mucus height we observed upon short term exposure to PGE2 is largely due to ion and fluid secretion-induced swelling of pre-existing mucus.

As the structural integrity of the colonic mucus layer is an essential part of the intestinal barrier, local changes in mucin organization or concentration due to swelling could affect structural properties of the mucus as well. Notably, the inner impenetrable mucus layer was preserved during PGE2-induced swelling of the mucus layer (Figure 4G, Supplementary Figure 8). Furthermore, when we exposed the mucus layers of control and PGE2-treated Colon Chips to increasing flow velocities, we found that the mucus layers in both chips exhibited similar bending angles, and hence, similar material responses to shear stress (Supplementary Figure 9, **Supplementary Video 1**). This suggests that despite changes in hydration state of the mucus layer, its gross structural integrity was maintained.

### DISCUSSION

Although the structure and function of the human colonic mucus bilayer is highly relevant for intestinal pathophysiology, previous investigation of its properties could only be carried in short-term (< 1 day) *ex vivo* tissue explants. This is because cultured human colonic epithelial cells do not produce a thick mucus layer with a normal bilayer structure in organoids, TW cultures, or any other *in vitr*o model ^17^. In contrast, in the present study, we showed that a microfluidic 2-channel human Colon Chip enables long-term culture of primary human colonic epithelial cells under dynamic flow conditions. Moreover, this system supports the spontaneous differentiation of large numbers of mucus-producing goblet cells at similar levels to those observed in human colon *in vivo*, while still maintaining a healthy subpopulation of proliferative cells. Importantly, under these culture conditions, the human colonic epithelial cells produced a mucus bilayer containing an impenetrable layer closely apposed to the apical surface of the epithelium, directly overlaid by a penetrable mucus layer, with a total thickness of 500-600 μm, which is similar to that seen in living human colon *in vivo* ^20^. Thus, this is the first culture method to recapitulate the development of a thick human colonic mucus layer with its unique bilayer structure.

Another novel feature of the human Colon Chip method is that the optical clarity of the microfluidic device allows live non-invasive visual analysis of mucus accumulation and physiology over time in culture. The dynamic changes in mucus layer thickness induced *in vivo* by the inflammatory mediator, PGE2, could be replicated, quantified, and analyzed on-chip. This revealed that rapid changes in mucus layer height after short term exposure to PGE2 are primarily mediated by altering the hydration state of pre-existing mucus via ion secretion through NKCC1, and not due to additional mucus secretion. Furthermore, our data indicate that colonic mucus may be able to undergo significant expansion without losing barrier-function or structural stability, highlighting the remarkable characteristics of this physiologically important structure. These findings demonstrate the usefulness of the Colon Chip as an *in vitro* tool for evaluation of mucus structure and function, which could advance our understanding of mucus physiology in disease contexts. Considering recent advances in bacterial co-cultures in intestinal microfluidic models ^23, 24, 35, 36^, this microfluidic Colon Chip lined by patient-derived colonic epithelial cells may also facilitate development of new therapeutics or probiotics that modulate the mucus barrier, as well as provide a novel testbed for personalized medicine.

## SYNOPSIS

An in vitro method is described for studying colonic mucus physiology by integrating primary human colonic epithelial cells in a microfluidic organ-on-a-chip device. The Colon Chip produces a mucus layer with thickness and bilayered microstructure similar to the human colon.

## AUTHOR CONTRIBUTIONS

A.S.-P. designed, executed and analyzed all experiments, with input and supervision from D.E.I., D.B.C., R.P.-B, and O.L.; D.B.C. helped with flow cytometry; D.B.C., A.T. and V.F. assisted with data interpretation; C.A.R. and D.T.B. provided advice on establishment of organoid cultures and interpretation of data; T.C.F. assisted with fluorescence and transmitted light microscopy; T. D. helped with performance of chip experiments and analysis of bead data; C.F. assisted with the alcian blue assay; V.F. sectioned the TWs; S.J.-F. and J.W. helped with scanning electron microscopy; M.K. and A.T. helped with develop the Colon Chip method; D.B.C., A.T., M.K., helped with generation of human organoid cultures from resections. E.S. generated the human organoid cultures from biopsies; A.S.-P. and D.E.I. wrote the manuscript with input from all co-authors.

## ABBREVIATIONS

Colon Chip: Colon-on-a-Chip
CFTR: Cystic fibrosis transmembrane conductance regulator
DF: Dark Field
MUC2: Mucin 2
Organ Chip: Organ-on-a-Chip
PDMS: Polydimethylsiloxane
PGE2: Prostaglandin E2
TW: Transwell
UC: Ulcerative Colitis

## ACKNOWLEDGEMENT

We thank A. Nordstrom (Harvard Medical School Electron Microscopy Facility) for help with scanning electron microscopy, G. Phelps for assistance with mucus layer analysis, P. Wallisch, C. Ng, and B. Calmari for technical assistance, and A. Chalkiadaki and A. Monreal for their guidance and excellent advice.

## MATERIALS AND METHODS

### Isolation of human colon epithelial cells

Human colon epithelium was isolated from colon resections or endoscopic tissue biopsies. Full thickness pieces of the human colon were obtained anonymously from healthy regions of colonic resection specimens processed in the Department of Pathology at Massachusetts General Hospital under an existing Institutional Review Board approved protocol (#2015P001859). Specimens were restricted to those with (non-neoplastic) disorders, and regions collected for isolation were determined to be healthy based on gross examination. Endoscopic biopsies were collected from macroscopically grossly unaffected regions of the colon of de-identified pediatric and young adult patients undergoing endoscopy for abdominal complaints. Informed consent and developmentally-appropriate assent were obtained at Boston Children’s Hospital from the donors’ guardian and the donor, respectively. All methods were carried out in accordance with the Institutional Review Board of Boston Children’s Hospital (Protocol number IRB-P00000529) approval.

For the isolation of colonic crypts, colon resections were processed by removing the colon epithelium with lamina propria, and then the epithelial layer or the entire biopsy was digested with 2 mg ml^−1^ collagenase I (Thermo Fisher Scientific, 17100-017) supplemented with 10 μM Y-27632 (Y0503, Sigma-Aldrich) for 40 min at 37 °C with intermitted agitation, as described^22, 37^. Colon organoids were grown embedded in growth factor-reduced Matrigel (354230, Lot 7317015, Corning) and stem cell expansion medium supplemented with 10 μM Y-27632 ^19, 22^. Stem cell expansion medium is composed of advanced DMEM F12 (12634-010, Thermo Fisher Scientific) supplemented with: L-WRN (Wnt3a, R-spondin, noggin) conditioned medium (65% vol vol^-1^) (produced by the CRL-3276 cell line, ATCC), 1x GlutaMAX (35050-061, Thermo Fisher Scientific), 10mM HEPES (15630-106, Thermo Fisher Scientific), recombinant murine epidermal growth factor (50 ng ml^-1^) (315-09, Peprotech), 1x N2 supplement (17502-048, Thermo Fisher Scientific), 1x B27 supplement (17504-044, Thermo Fisher Scientific), 10 nM human [Leu15]-gastrin I (G9145, Sigma-Aldrich), 1 mM n-acetyl cysteine (A5099, Sigma-Aldrich), 10 mM nicotinamide (N0636, Sigma-Aldrich), 10 μM SB202190 (S7067, Sigma-Aldrich), 500 nM A83-01 (2939, Tocris), and primocin (100 μg ml^-1^) (ant-pm-1, InvivoGen).

### Colon Chip Cultures

The Organ Chips composed of poly-dimethylsiloxane (PDMS) and containing two parallel microchannels (apical channel 1000 x 1000 μm and basal channel 1000 x 200 μm; width x height) separated by a 50 μm thick PDMS porous membrane (7 μm pore diameter, 40 μm spacing) were purchased from Emulate Inc. (RE00001024 Basic Research Kit, Emulate Inc.). After activation of the channel surfaces with 0.5 mg ml^-1^ sulfo-SANPAH solution (A35395, Thermo Fisher Scientific), the inner surfaces of both channels and the porous PDMS membrane were coated with 200 µg ml^−1^ rat tail collagen type I (354236, Corning) and 1 % Matrigel (354230, Lot 7317015, Corning) in Dulbecco’s phosphate-buffered saline (DPBS), as previously described^22, 24^. Colon organoids were then isolated from Matrigel by incubating in cell recovery buffer (354253, BD) for 40 min on ice and the spun down at 400g for 5 minutes at 4°C. The colonic organoids were fragmented by incubating them in TrypLE™ Express Enzyme (12605010, Thermo Fisher Scientific) diluted in DPBS 1:1 (vol:vol) supplemented with 10 μM Y-27632 (2 mL/ well of a 24 well plate) for 1 min 45 seconds in a 37°C water bath. After adding the same volume of stem cell expansion medium with 10 μM Y-27632, the organoids fragments were spun down at 400g for 5 min at 4°C and then resuspended at 6 x 10^6^ cells ml^-1^. The colonic organoid fragments were seeded on the ECM-coated membrane in the apical channel of the Colon Chip (6 x 10^5^ cells cm^-2^) in stem cell expansion medium supplemented with 10 μM Y-27632 while filling the basal channel with the same medium, and the chips were incubated overnight at 37°C under 5% CO2 to promote cell adhesion. The following day both channels were washed once with stem cell expansion medium, and then the chips were inserted into Pod™ portable modules (RE00001024 Basic Research Kit, Emulate Inc.) and placed within a ZOË™ culture instrument (Emulate Inc.), where they were perfused (60 μl h^−1^) with stem cell expansion medium in the basal channel and HBSS with calcium and magnesium (21-023-cv, Corning) supplemented with 100 μg ml^-1^ primocin (ant-pm-1, InvivoGen) in the apical channel.

### Transwell insert cultures

Transwell® culture inserts (TWs; 6.5 mm) with 0.4 µm pore polyester membrane (3470, Corning) were coated as described above for the chip, and seeded with colon organoids fragments at the same density (6 x 10^5^ cells cm^-2^) on the top side of the TWs in stem cell expansion medium supplemented with 10 μM Y-27632, and the same medium was added to the bottom chamber. As with the chips, the TWs were incubated overnight at 37°C under 5% CO_2_, and the following day, the TW was washed once with stem cell expansion medium before adding 1 ml stem cell expansion medium on the basal side and 250 μl of HBSS with calcium and magnesium supplemented with 100 μg ml^-1^ primocin to the apical side; medium was changed every 2 days thereafter.

### Organoid cultures

Colon organoid fragments were resuspended in growth factor-reduced Matrigel at 1 x 10^6^ cells ml^-1^ and plated in 24 or 48 well plates (50 or 10 μl drops/well, respectively), and covered with stem cell expansion medium supplemented with 10 μM Y-27632 (500 μl or 200 μl/ well, respectively). Stem cell expansion medium was changed every 2 days thereafter.

### Immunofluorescent microscopy

Colon Chip were fixed with 200 μl of 2% paraformaldehyde (PFA) (Electron Microscopy Sciences, 15730), 25mM HEPES (Thermo Fisher Scientific, 15630-080) in DPBS (Gibco, 14190-144) with 200 μl filter tips at 4°C on a rocker overnight. TWs and organoids were fixed at room temperature for 15 min. Chips were either stained directly or sectioned at 250 μm with a vibratome (VT1000S, Leica). Fresh frozen 7 µm *in vivo* tissue sections were fixed with 2% PFA for 12 min at 4°C. All samples were blocked and permeabilized using 0.1% Triton X-100 (X100, Sigma), 5% BSA (A2153, Sigma) in DPBS for 1 h at room temperature. Samples were then stained overnight at 4°C in 2% BSA in DPBS with primary antibodies: anti-MUC2 (H-9, sc-515106, Santa Cruz, 1:100), anti-E-Cadherin (HECD-1, ab1416, abcam, 1:100), anti-ZO-1 (ZO1-1A12, 33-9100, abcam, 1:200). The next day, after 3 washes of PBS, samples were then stained overnight at 4°C in 2% BSA in DPBS containing secondary antibodies and phalloidin: Goat anti-mouse-IgG1 Alexa Fluor 647 (A-21240, Invitrogen, 1:100), goat anti-mouse IgG2b Alexa Fluor 555 (A-21147, Invitrogen, 1:500), Phalloidin Alexa Fluor 488 (A12379, Invitrogen, 1:200), Phalloidin Alexa Fluor 647 (A22287, Invitrogen, 1:200). The next day, after 3 DPBS washes, staining with Hoechst 33342 (H3570, Life Technologies, 1:2,000) for 30 min was performed.

Images were taken using a Leica SP5 laser scanning confocal immunofluorescence microscope with a 680-1080nm multiphoton pulsed IR laser Chameleon Vision 2 with pre compensation and Non-Descanned Detectors, a 470-670nm nm white light laser, 488 nm argon laser and coupled to HyD detectors. Acquired images were analyzed using IMARIS software (Bitplane).

### Flow Cytometry

For assessment of proliferating cells, culture medium (stem cell expansion medium basally, HBSS apically) containing 10 µM EdU (Click-iT™ Plus EdU Alexa Fluor™ 350 Flow Cytometry Assay Kit, C10645, Invitrogen) was perfused through the apical and basal channels of the chip for 18h prior to cell harvest. Cells were isolated from the Colon Chips and TWs by incubation in 1mg ml^-1^ collagenase IV in TrypLE supplemented with 10 μM Y-27632 (100 μL/ channel in the chips; 100 μL above and 500 μL below the membrane in TWs) for 1h at 37°C. Detached cell fragments were incubated for an addition 45 min at 37°C up to 1h until a single cell suspension was obtained. Epithelial cells were isolated from colonic organoids as described above except that the organoids, after extraction from Matrigel, were incubated in 200 μl enzyme solution for 45 min up to 1h at 37°C. Cells were also isolated from human colon resections by dissecting the tissue as described above. Colonic crypt isolation and digestion into single cells was performed as previously described^38^. In short, minced tissue was incubated in 8mM EDTA in DPBS (14190-144, Gibco) while slowly rotated for 75 min at 4°C, followed by vigorous shaking of the sample to enrich for dissociated colonic crypts. To obtain a single cell suspension, colonic crypts were incubated in Disaggregation Medium (Advanced DMEM/F12, 1x GlutaMAX, 10 mM HEPES, 1x N-2, 1x B-27, 10 mM nicotinamide, 1 mM N-acetyl-L-cysteine, 10 µM Y-27632, 2U ml^-1^ dispase (17105041, Gibco), 200 KU DNAse I ml^-1^ (D5025, Sigma) and incubated for 30 min at 37°C with occasional agitation. All harvested cells were centrifuged, resuspended in flow staining buffer composed of 1% FBS (Gibco, 10082-147), 25mM HEPES (15630-080, Thermo Fisher Scientific), 1mM EDTA (15575-020, Thermo Fisher Scientific), and 0.05% sodium azide (BDH7465-2, VWR) in DPBS (14190-144, Gibco). Surface staining was performed in 100 ul staining buffer for 30 min, followed by fixation in 2% PFA (panel 1 and 3) for 15 min or overnight fixation with eBioscience™ Foxp3 / Transcription Factor Staining Buffer Set (00-5523-00, Invitrogen) (panel 2). After fixation, EdU staining was performed following manufactures instructions (Click-iT™ Plus EdU Alexa Fluor™ 350 Flow Cytometry Assay Kit, C10645, Invitrogen), followed by intracellular staining in 1x saponin (Click-iT™ Plus EdU Alexa Fluor™ 350 Flow Cytometry Assay Kit, C10645, Invitrogen).

Panel 1 (2% PFA fixation): anti-MUC2 (H-9, sc-515106, Santa Cruz, 1:100), anti-mouse-IgG2b-546 (A-21143, Invitrogen, dilution 1:100), Click-iT™ Plus EdU Alexa Fluor™ 350 Flow Cytometry Assay Kit (C10645, Invitrogen).

Panel 2 (Foxp3): anti-Ki67 (Ki-67, 350514, BioLegend, dilution 1:20), Click-iT™ Plus EdU Alexa Fluor™ 350 Flow Cytometry Assay Kit (C10645, Invitrogen).

Panel 3 (fresh *in vivo* tissue): anti-CD45-Brilliant Violet 570 (HI30 clone, 304034, BioLegend, dilution 1:50), anti-CD235a-Pacific Blue (HI264 clone, 349108, BioLegend, dilution 1:40), anti-CD11b-Brilliant Violet 570 (clone, cat, BioLegend, dilution 1:40), anti-CD31-421 (clone, cat, BioLegend, dilution 1:20), EpCAM-PE/Cy7 (CO17-1A, cat, BioLegend, dilution 1:20), anti-MUC2 (H-9, sc-515106, Santa Cruz, 1:100), anti-mouse-IgG2b Alexa Fluor 546 (A-21143, Invitrogen, dilution 1:100).

All panels included 20 nM Syto16 (S7578, Thermo Fisher Scientific, dilution: 1:500), Zombie NIR dye (423106, BioLegend, dilution: 1:500), Fc Block (BioLegend, 422302, dilution 1:20). Stained cells were analyzed using the LSRFortessa (BD Biosciences). Results were analyzed using FlowJo V10 software (Flowjo, LLC).

### Permeability

Cascade Blue™ hydrazide Trilithium Salt (550 Da) (C3239, Invitrogen) at 50 µg ml^-1^ in HBSS was added to the top epithelial channel of the Colon Chip to assess barrier permeability. The concentration of dye that diffused through the membrane into basal channel was measured in the effluent, and apparent paracellular permeability (Papp) was calculated as previously described^22^.

### Light microscopy

The top view Colon Chip images were acquired using a differential interface contrast (DIC) or phase contrast microscopy (Zeiss Axio Observer Z1). Frozen sections were obtained during different stages of the isolation of colon epithelial cells, stained for H&E and imaged.

### Side view imaging of mucus accumulation on-chip

To image mucus accumulation in living cultures on-chip, ∼2 mm of PDMS were cut away from each side of the chip parallel to the channels using a razor blade secured in a press. The chips were rotated onto one side on a glass slide coated with Glycerine-Solution (11513872, Leica) and the top side was covered with glycerin and a cover glass. The images were acquired with an inverted microscope (Zeiss Axio Observer Z1) using a 2.5x objective (0.06 NA, 441010-9901, Zeiss) and condenser (0.35 NA, 424241-0000-000, Zeiss) with phase ring 2 used for DF imaging. Fluorescent images were acquired using an X-cite LED light source (Excelitas Technologies). Mucus height and area were analyzed in side view images of the Colon Chip using Fiji software.

### PGE2 treatment

For long-term PGE2 studies, the basal channel of the Colon Chips was perfused with stem cell expansion medium 1.4 nM PGE2 (P5640, Sigma) for 6 days, starting at day 2. To quantify cell proliferation, 10 µM EdU (C10645, Invitrogen) was perfused through both channels of the chip for 18 hours prior to enzymatically detaching cells on day 8 and carry out flow cytometric analysis. Short term treatment with PGE2 and ion channel inhibitors was performed at 7 days after monolayer formation. In short, Colon Chips were perfused with medium (stem cell expansion medium basally, HBSS apically) containing 50 µM CFTRinh-172 (S7139, Selleckchem), 20 µM XE-991 Dihydrochloride (20010, tocris), 100 µM Bumetanide (S1287, Selleckchem), or combination of the 3 inhibitors at 60 µl h^-1^ for 4h, followed by side view imaging to determine the baseline height of the mucus layer prior to PGE2 treatment. After baseline side view imaging, the chips were then switched to co-treatment with 1.4 nM PGE2 (basal channel) and the respective ion channel inhibitors (apically and basally). After 4h of co-treatment with inhibitors and PGE2, side view imaging was performed to determine the swelling of the mucus layer.

### Scanning Electron Microscopy

To visualize the epithelium and mucus layer on-chip, SEM analysis was carried out using Colon Chips that had a top channel that was not irreversibly bonded to the membrane, which allowed the device to be dismantled manually as describe previously^39^. Cells were fixed with 4% PFA (157–4, Electron Microscopy Sciences) and 2.5% glutaraldehyde (G7776; Sigma) in DPBS and incubated in 0.5% osmium tetroxide (19152, Electron Microscopy Sciences) in 0.1 M sodium cacodylate buffer (pH 7.4) before serial dehydration in ethanol. Samples were then dried using a critical point drier and imaged using field emission SEM (Hitachi S-4700).

For cross sectional SEM images, Colon Chips were fixed with 2.5% Glutaraldehyde in 0.1 M Phosphate Buffer (P5244, Sigma) overnight and washed with water before being flash-frozen in liquid nitrogen. Frozen chips were sectioned into 5 mm cross sections on dry ice, lyophilized, mounted on aluminum pin mounts with conductive carbon tape, sputter-coated with gold, and examined with a Tescan Vega3 GMU scanning electron microscope.

### Analysis of inner and outer mucus layer

Cells in the Colon Chips were stained for live cells by perfusing both channel for 30 min with medium (stem cell expansion medium basally, HBSS apically) containing 10 µM Calcein AM (C3100MP, Invitrogen) (200 µl h^-1^); the medium perfused through the apical channel also contained 1 µm FluoSpheres™ Carboxylate-Modified Microspheres (70 µl ml^-1^; F13083, Invitrogen). Colon Chips were incubated under static conditions for 40 min to allow the fluorescent beads to settle, and then z-stack images of 2-3 areas of each chip were collected using a Leica SP5 confocal microscope. Calcein AM was imaged with a 488 nm Argon laser, and beads were visualized using the multiphoton laser at 1000 nm. Confocal z-stacks were reconstructed and analyzed using IMARIS software (Bitplane). To determine the thicknesses of the inner and outer mucus layers, a Gaussian distribution was fit to the data using Matlab and height of the outer layer was determined using the middle 90% of the Gaussian distribution of the beads. The inner layer was set as the distance between the apical cell surface and the lower bound of the outer layer.

### Shear stress deformation assay

Increasing flow rates were applied to the Colon Chips using a Fusion Touch Syringe Pump. Side view images were acquired by transmitted light microscopy and movies were generated during flow and stop cycles of the pump at 1.6 ml h^-1^, 6 ml h^-1^, 10 ml h^-1^. Images were then analyzed using Fiji software by tracing movement of mucus strains while flow was applied compared to the final position after flow was stopped. Angle between flow and no flow was calculated (great angle equals great deformation).

### Drawings

All drawings were created with BioRender.

### Alciann blue mucin assay

Mucus was loosened from the apical surfaces of the colon chips by reducing disulfide bonds with 250 mM Tris (2-carboxyethyl) phosphine (C4706, Sigma-Aldrich). After 1h incubation, the mucus layer was mechanically separated by washing the apical channel with phosphate buffered saline (PBS). Samples were frozen, lyophilized overnight and reconstituted in PBS. Total mucous amount was determined using an alcian blue colorimetric assay adapted from Hall et al (1980)^40^. Briefly, a standard curve was created using serial dilutions of submaxillary gland mucin (Sigma) ranging from 0 μg ml^-1^ to 500 μg mL^-1^. Samples were diluted into the linear range of the curve using PBS. Samples and standards were equilibrated with filtered Richard Allan Scientific^TM^ alcian blue (88043, Thermo Fisher Scientific) for 2h. The resulting precipitant was separated by centrifugation at 1870 g for 30 min. This was followed by a series of wash/spin cycles at 1870 g in a resuspension buffer composed of 40% ethanol, 0.1 M acetic acid and 25 mM magnesium chloride. The mucin pellets were then dissociated with a 10% SDS solution (71736, Sigma) and absorbance was measure with a microplate reader (Synergy HT, BioTek) at 620 nm. Mucin concentration values for samples were interpolated from a linear fit of the standard curve.

### Statistical Analysis

All graphs are depicted as mean ± standard error of the mean (SEM) and significant differences between two groups were determined using two-tailed unpaired Student’s t-test. To determine significant differences between 3 groups or more, one-way ANOVA with Tukey’s multiple comparisons test was used. With 3 groups or more and 2 independent variables, 2ways ANOVA with Tukey’s multiple comparisons was used to determine statistical significance. Prism 7 (GraphPad Software) was used for statistical analysis.

### Data Availability

The authors declare that the data supporting the findings of this study are available within the article and its supplementary information files.

All authors had access to the study data and had reviewed and approved the final manuscript.

## SUPPLEMENTARY FIGURE LEGENDS

**Supplementary Figure 1:**
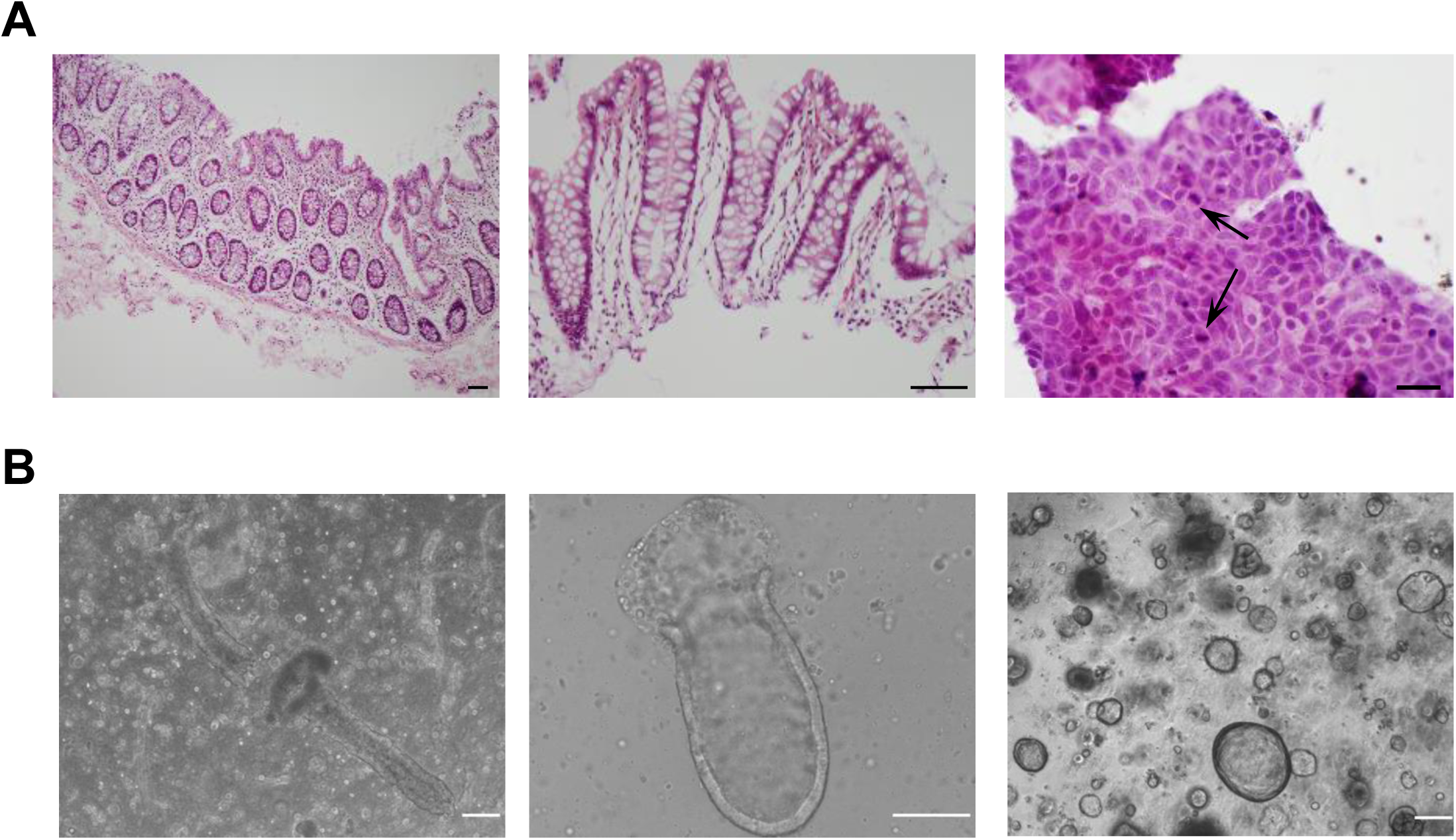
Establishment of primary colon organoids from human colon resections. (A) H&E staining of human colon *in vivo* sections (6 µm). Colon tissue with intact epithelium, lamina propria, muscularis mucosa and submucosa (left, bar, 100 µm). Dissociated epithelium including lamina propria (middle, bar, 100 µm). Smear of a 2-day colon organoid in matrigel culture (right, bar, 20 µm). The arrows are marking mitotically active epithelial cells. (B) Primary human colonic epithelium in matrigel cultures. Colonic crypts embedded in Matrigel after isolation from human colon resection (left). Shortening of crypt structure embedded in Matrigel one day after isolation (middle). Closed colonic crypts (colon organoids) in Matrigel cultures 7 days after isolation (right) (bars, 200 µm).

**Supplementary Figure 2:**
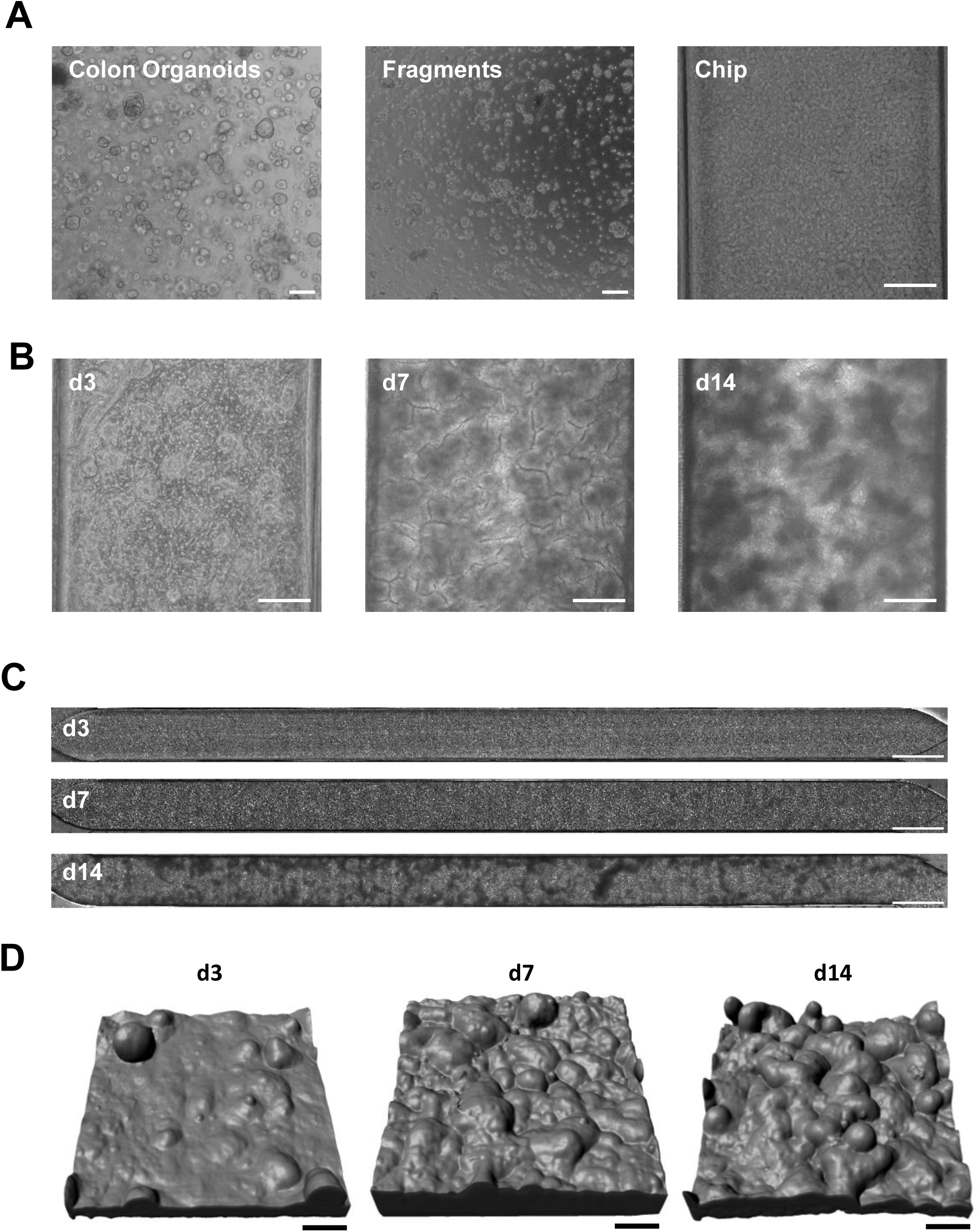
Development of the Colon Chip. (A) Phase contrast images of the preparation of primary human colonic epithelial cells for seeding on the Chip starting with colon organoids that are fragmented before seeding on chip (bars, 200 µm). (B) Development of epithelial cells on Colon Chip over 14 days after monolayer formation with phase contrast images (bars, 200 µm). (C) Full length DIC images of the epithelial cells cultured on the Colon Chip over 14 days after monolayer formation (bars, 1000 µm). (D) 3D reconstruction of confocal microscopy z-stack images of the epithelial cells on Colon Chip based on F-actin staining on day 3, day 7 and day 14 of culture after monolayer formation (bars, 100 µm).

**Supplementary Figure 3:**
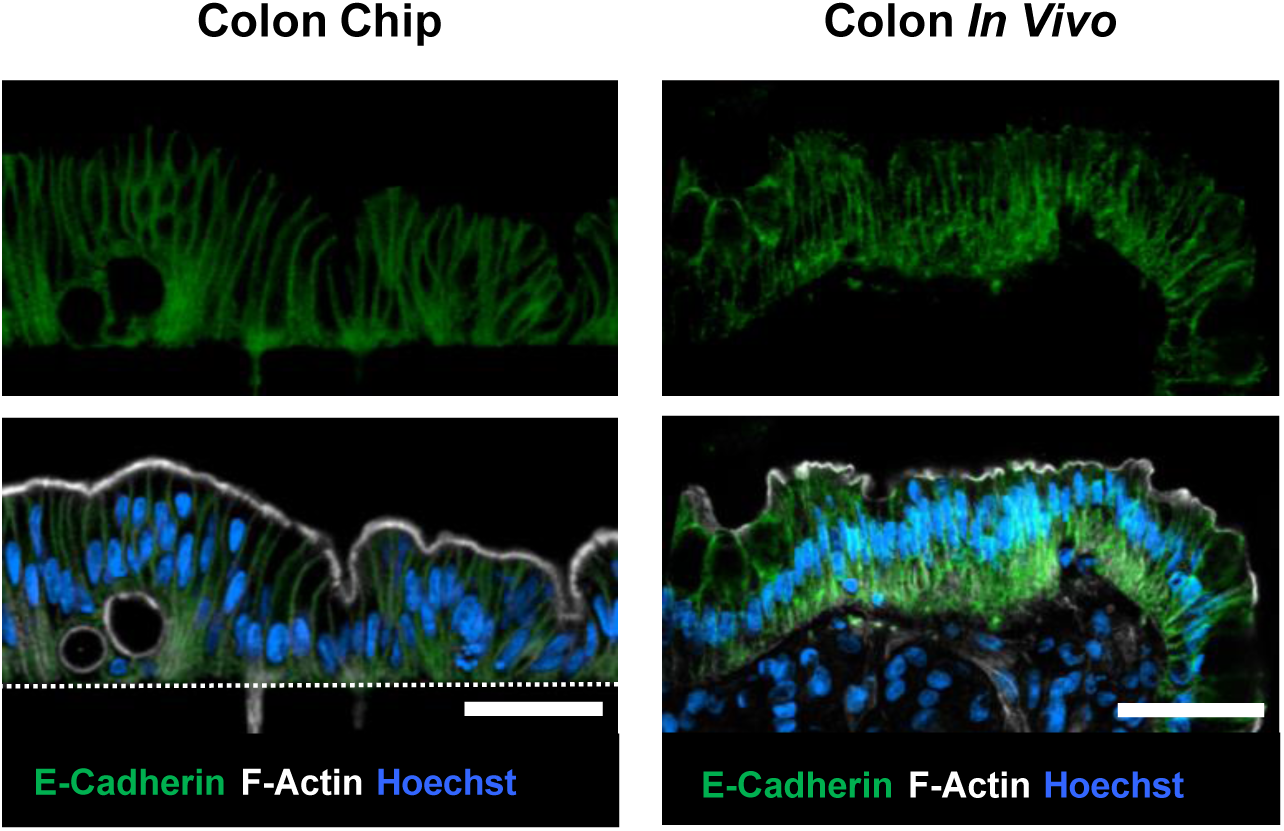
Formation of a polarized epithelium within the Colon Chip. Cross-sectional, immunofluorescence confocal microscopic images of the Colon Chip compared with human colon *in vivo* showing a polarized epithelium with adherens junctions labeled with E-Cadherin (green) and brush border stained for F-actin (gray) restricted to the apical regions, and Hoechst-stained nuclei (blue) localized at the cell base (bar, 50 µm; white dashed line, top of the porous PDMS membrane in the Colon Chip).

**Supplementary Figure 4:**
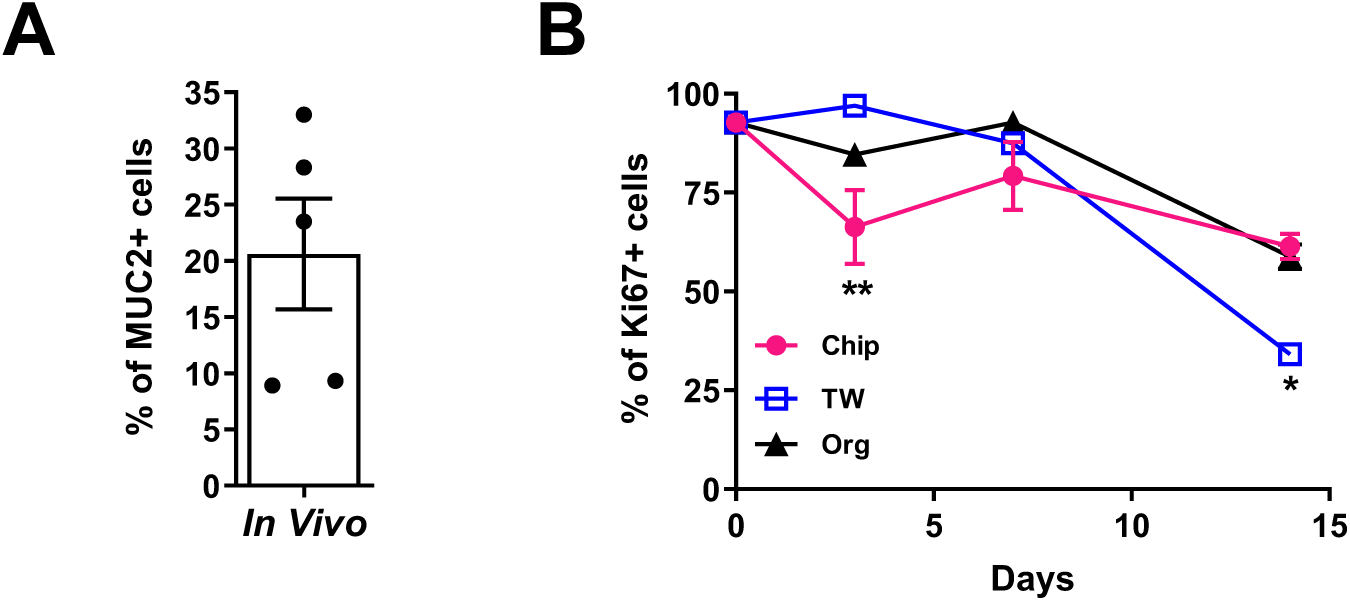
In Vivo colon goblet cell quantification. (A) Quantification of MUC2+ cells by flow cytometry on cells isolated from human colon tissue (n=2 donors, 2-3 samples per donor. (B) Quantification of proliferating cells by Ki67 from Colon Chip (Chip), TWs and colon organoids (Org) on day 3, day 7 and day 14 by flow cytometry (n=3-8 devices, 2 experiments compiled; ** *p*<0.01). (all data represented as mean ± SEM).

**Supplementary Figure 5:**
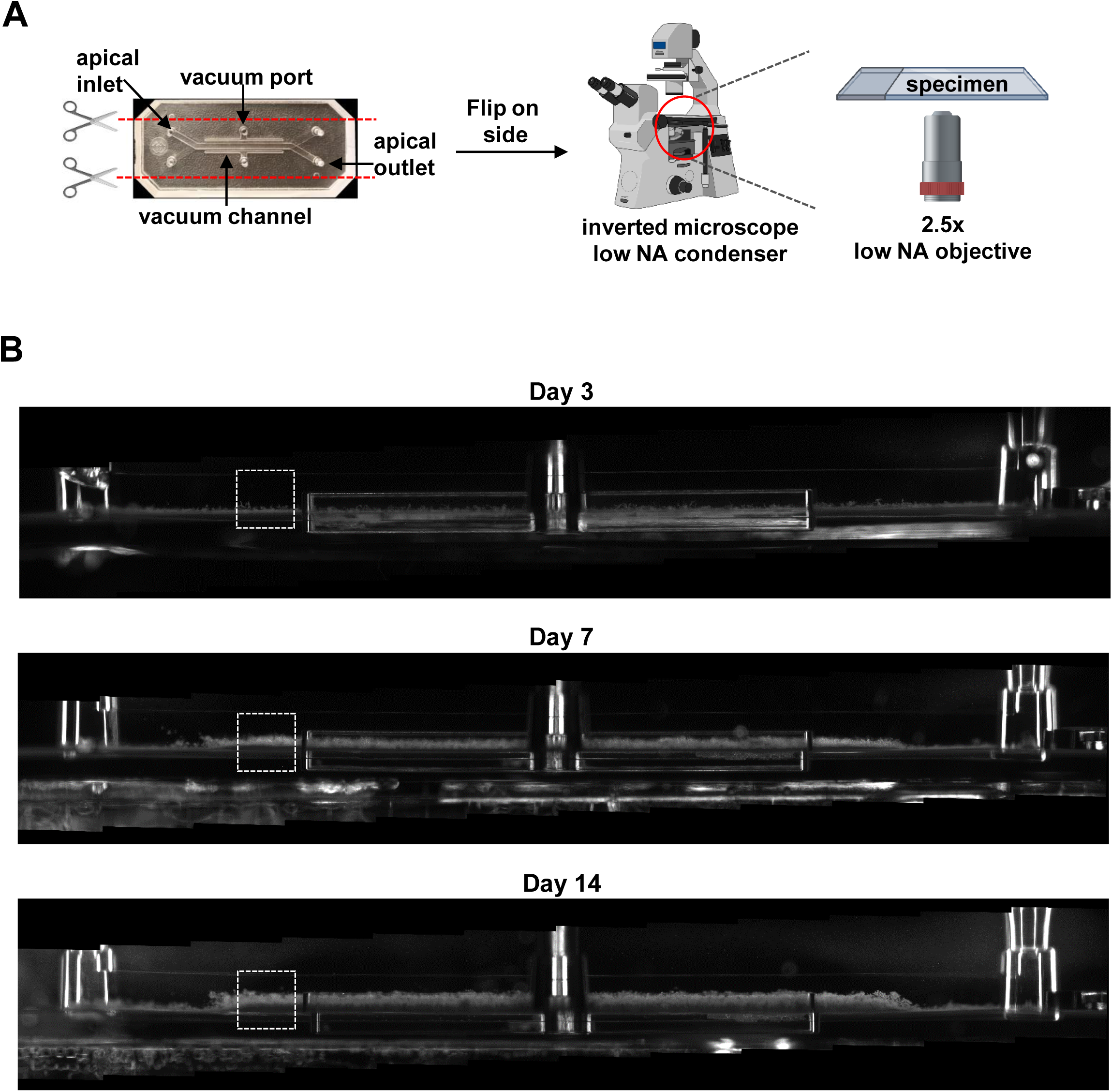
Visualization of mucus layer height on-chip using Dark Field Microscopy. (A) Schematic of preparation of chip for side view imaging. (B) Whole chip side view images live of Colon Chips at day 3, 7 and 14 by dark field (DF) microscopy showing cells and mucus growth over time with dashed square indicating regions of the chips displayed in Figure 3C.

**Supplementary Figure 6:**
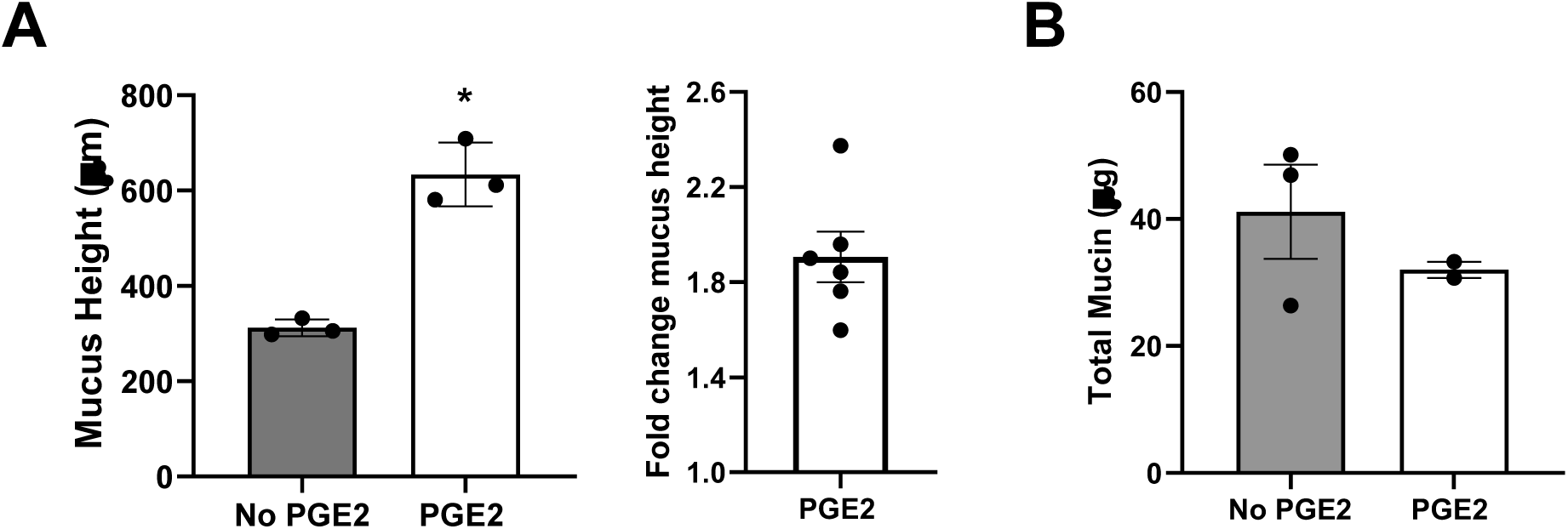
PGE2-induced mucus swelling. (A) Change in mucus height after 4h PGE2 treatment over the mucus height before treatment in the same chips (n=6, 2 experiments compiled). (B) Total mucin amount per chip with and without 4h PGE2 (n=2-3) not statistically significant by t-test. (all data represent mean ± SEM).

**Supplementary Figure 7:**
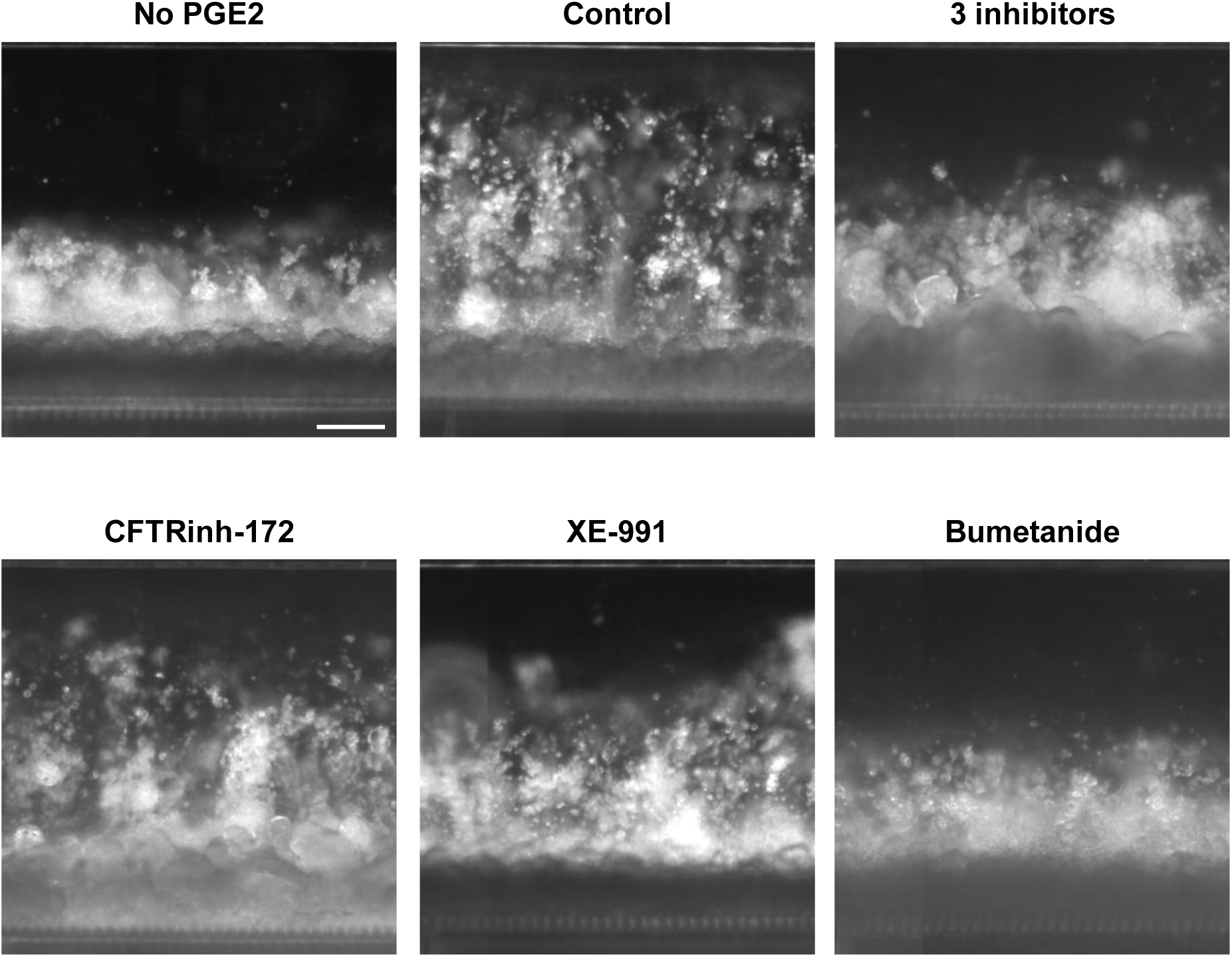
Modulation of PGE2-induced changes in mucus layer thickness by ion channel inhibition. DF side view images of a Colon Chip with epithelial cells and mucus layer before (No PGE2) and after PGE2 treatment for 4 hours (Control) or with CFTRinh-172 (50 µM), XE-991 dihydrochloride (20 µM), bumetanide (100 µM) or a combination of all three ion channel inhibitors (bar, 200 µm).

**Supplementary Figure 8:**
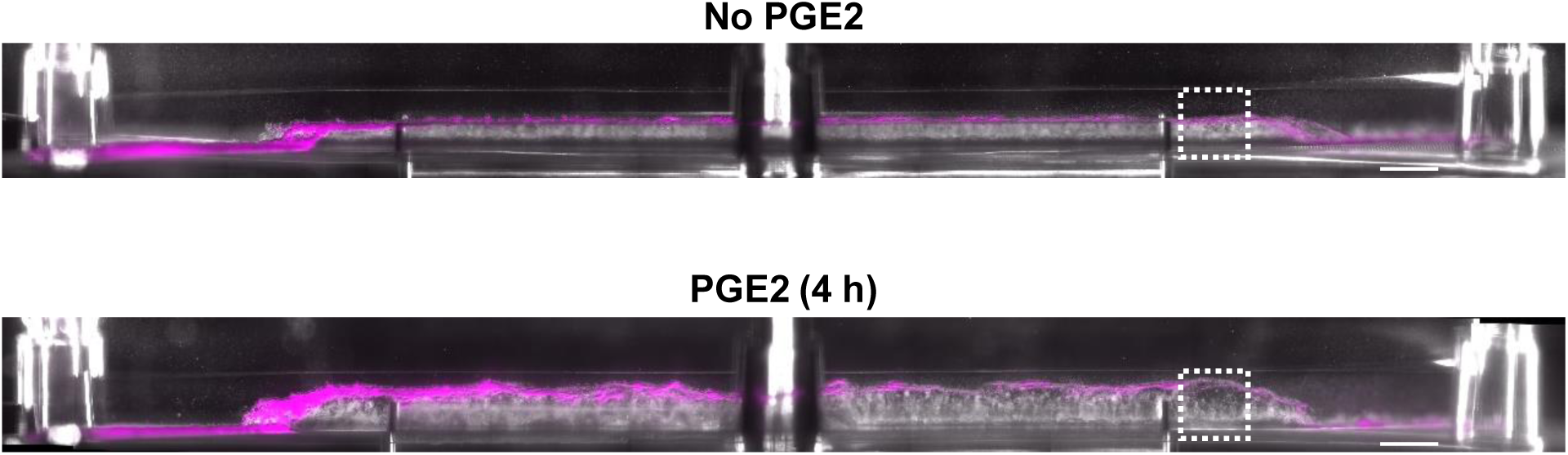
Inner mucus layer is not affected by PGE2. Side view DF and fluorescent image of chip with mucus layer and 1 µm fluorescent beads (magenta) without and with PGE2 treatment with dashed square indicating a region of the chips represented in the higher magnification images in Figure 4G (bars, 1000 µm).

**Supplementary Figure 9:**
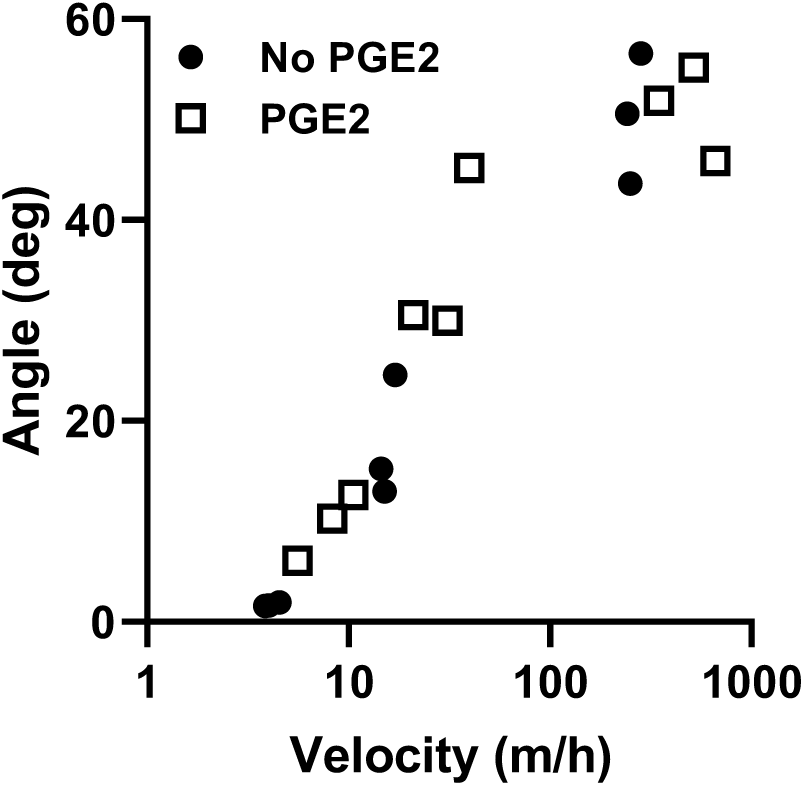
Response of mucus layer to shear stress. Response of the mucus layer to shear forces measured in angle over increasing velocity, with a higher angle being equivalent to stronger bending of the mucus layer (each data point represents 1 chip).

**Supplementary Video 1: Shear force assay.** Side view images compiled to movies at 6 ml h-1 and 10 ml h-1 flow rate without (No PGE2) or with 4h of PGE2 treatment (4h PGE2).

